# Bioinformatic and cell-based tools for pooled CRISPR knockout screening in mosquitos

**DOI:** 10.1101/2021.03.29.437496

**Authors:** Raghuvir Viswanatha, Enzo Mameli, Jonathan Rodiger, Pierre Merckaert, Fabiana Feitosa-Suntheimer, Tonya M. Colpitts, Stephanie E. Mohr, Yanhui Hu, Norbert Perrimon

**Affiliations:** Department of Genetics, Blavatnik Institute, Harvard Medical School, Boston, MA 02115, USA; Department of Microbiology, National Emerging Infectious Diseases Laboratories, Boston University School of Medicine, 620 Albany Street, Boston, MA 02118, USA; HHMI, Harvard Medical School, Boston, MA 02115, USA

## Abstract

Mosquito-borne diseases present a worldwide public health burden. Genome-scale screening tools that could inform our understanding of mosquitos and their control are lacking. Here, we adapt a recombination-mediated cassette exchange system for delivery of CRISPR sgRNA libraries into cell lines from several mosquito species and perform pooled CRISPR screens in an *Anopheles* cell line. To implement this method, we engineered modified mosquito cell lines, validated promoters and developed bioinformatics tools for multiple mosquito species.

---

Mosquito-borne diseases include a vast repertoire of viral, bacterial and parasitic diseases of medical and veterinary importance, with malaria alone causing nearly half a million human deaths each year^1^. Current efforts to fight malaria and other mosquito-transmitted diseases such as Dengue, Zika, Chikungunya and West Nile Virus rely on control of vector populations, mostly by means of insecticides. These measures are hampered by ever-increasing insecticide resistance. Alternative strategies under current development include those based on the use of endosymbiotic bacteria or gene-drives to suppress wild mosquito populations or replace them with disease-refractory mosquitos^2^.

Studies of mosquito-borne diseases would benefit of the availability of methods that allow large-scale functional cell-based screens, for example genome-scale screens for virus, bacteria or parasite entry, innate immunity, or resistance to insecticides and other toxins. Previously, we developed such a method in *Drosophila* cells, based on recombination-mediated cassette exchange (RMCE) system to deliver complex CRISPR sgRNA libraries. Here, we establish such an approach in mosquitos by generating engineered RMCE acceptor cell lines from multiple mosquito species, testing a series of promoter, and developing bioinformatic tools for CRISPR guide design and other applications. We demonstrate the robustness of the approach by performing a large-scale pooled CRISPR screen in the African malaria vector *Anopheles coluzzii*- derived cell line Sua-5B^3^.

To generate RMCE adapter cell lines as a platform for CRISPR screens, we first chose well characterized cell lines from three mosquito species that are susceptible to infection with biomedically important viruses or parasites, and for which genomic, transcriptomic, and small RNA sequencing data exist^4^: Sua-5B from *Anopheles coluzzii*^3^ (formerly *An. gambiae* M form), Hsu from *Culex quinquefasciatus*^5^, and C6/36 from *Aedes albopictus*^6^. Next, we applied MiMIC technology, which uses *Minos* transposition to deliver an RMCE acceptor, to add a docking site for recombination via the bacteriophage ΦC31 integrase^7,8^. This allows for stable integration of sgRNAs^9^. Modified cells are identified by the presence of an mCherry exon that becomes incorporated into a native gene. mCherry-expressing cells were isolated using fluorescence-activated cell sorting (FACS) and we selected a single, strongly mCherry-positive derivative cell-line from each parental line: Sua-5B-1E8 (*Anopheles*), Hsu-1.7 (*Culex*), and C6/36-HE8 (*Aedes*) (**Supplementary Figure 1**).

An incompletely addressed challenge for CRISPR genome engineering in mosquitos is the identity of optimal *pol III* promoters for heterologous expression of sgRNAs^10–14^. We performed side-by-side evaluations of eleven *pol III* promoters from four mosquito species, as well as a consensus sequence, in each of the three mosquito cell lines. To choose promoters, we first used BLAST and multiple alignment to identify orthologs of the *Drosophila* U6 promoter and chose eleven orthologous promoters from U6 snRNAs of *Anopheles, Culex*, or *Aedes* (**Fig. 1a**). When possible, we selected a minimum of three promoters per species, prioritizing U6 promoters for which RNA-seq data suggests they are expressed in cell lines and in adult tissues (see Methods). We confirmed that each mosquito U6 promoter included an intact *pol III* bipartite promoter motif. These were synthesized and inserted in the pLib6.4^9,15^ to generate a suite of vectors for expression of sgRNA under the control of different *pol III* promoters (**Supplementary Data 1**).

**Fig. 1.**
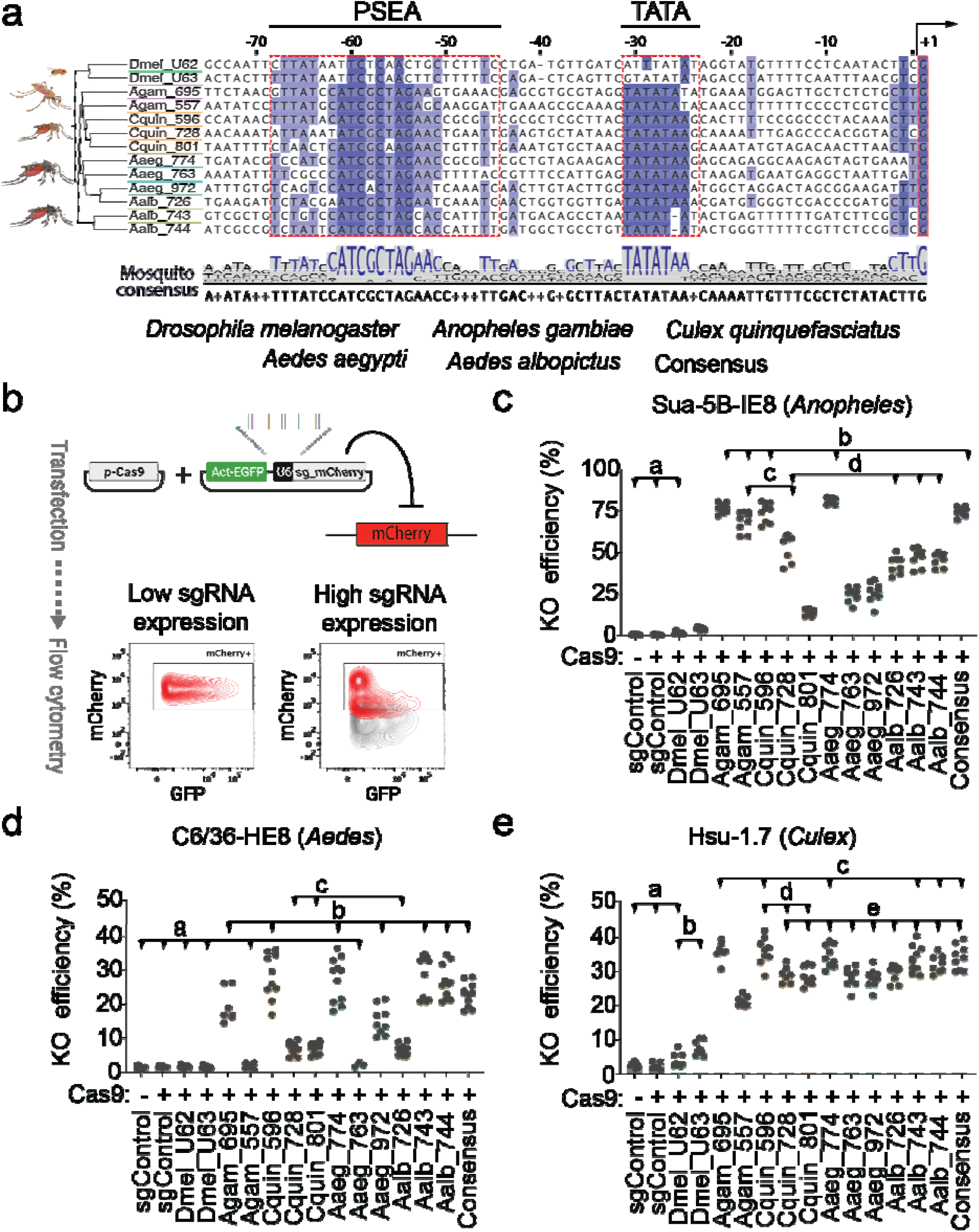
Identification of U6 promoters for sgRNA expression in mosquito cells and evaluation of CRISPR knockout efficiency. (**a**) Multiple alignment of selected U6 promoters highlighting the metazoan pol III promoter bipartite structure and a mosquito consensus sequence. The first transcribed base of the U6 snRNA, the TATA box and proximal sequence element A (PSEA) are boxed in red. The consensus sequence derived from mosquito sequences was used to design a synthetic U6 promoter. (**b**) Flow cytometry-based assay for evaluation of CRISPR knockout (KO) efficiency with different U6 promoters. Engineered mosquito cells expressing mCherry were co-transfected with a plasmid expressing Cas9 and a GFP reporter plasmid expressing the sgRNA targeting mCherry under the control of different U6 promoters. After transfection, cells were passaged for up to 12 days and then analyzed by flow cytometry. Only GFP^+^ cells were considered in the analysis. Relative KO efficiency was obtained calculating the ratio of mCherry-cells over total GFP transfected cells. (**c**) Evaluation of U6 promoter-specific CRISPR-KO efficiencies in three mosquito cell lines. Histogram bars represent the mean, dots represent the distribution of multiple replicates obtained from 3 independent experiments. Histogram colors denote the species of origin of the U6 promoters analyzed, shown with abbreviation of species name and three last letters of the corresponding Vectorbase gene ID. sgControl= pLib6.4-Agam_695 U6 expressing the empty BbsI cassette was used as control. Statistical analysis was performed using Brown-Forsythe ANOVA test followed by Dunnett’s multiple comparison test. Lowercase letter groupings denote differences not significant (P_Dunnett_ > 0.05). All differences between samples of different groupings are significant (P_Dunnett_ < 0.05).

Mosquito cells with genomically-encoded mCherry allowed us to use a flow cytometry-based dual reporter assay to directly compare knockout (KO) efficiency in cells expressing the same sgRNA from different *pol III* promoters (**Fig. 1b, Supplementary Figure 2a, Supplementary Data 2**). An advantage of this approach over a dual reporter assay using transfected targets is that integrated targets are expected to reproduce gene repair outcomes with the same dynamics as native genes. We used GFP as an indicator of the efficiency of co-transfection of the mCherry expressing cells with Cas9 and the newbuilt gRNA expressing vector targeting mCherry. The ratio of mCherry^-^ cells within the GFP transfected cells was then used to determine KO efficiency. In *Anopheles* cells, all mosquito promoters tested elicited measurable knockout, whereas *Drosophila* promoters failed (**Fig. 1c, Supplementary Figure 2b**). The native promoters (AGAP013695, AGAP013557) along with *Culex* CPIJ039596 and *Ae. aegypti* AAEL017774 showed the strongest activity, achieving approximately 75% KO efficiency relative to controls. In particular, AAEL017774 (mean=81.3 SD±1.9) and AGAP013695 (mean=76.6 SD±3) were the most efficient. The remaining promoters have moderate to low activity, and the mosquito consensus promoter performed similarly to the native promoters. In the *Culex* cell line, we observed a more uniform activity of mosquito U6 promoters, with an overall mean KO efficiency of about 30% (**Fig. 1d, Supplementary Figure 2b**). Notably, the results for CPIJ039596 obtained using this assay were slightly lower but overall comparable to CRISPR allele editing efficiency as verified by deep sequencing for the same promoter in our previous study^13^.

The most effective U6 promoters in *Ae. albopictus* C6/36-HE8 cells were the native promoters AALF029743-4 (mean=28.6 SD±6.1; mean=26.4 SD±4.9), *Culex* CPIJ039596 and *Ae. aegypti* AAEL017774, with about 27% mean KO efficiency (**Fig. 1d, Supplementary Figure 2b**). Interestingly, *Culex* CPIJ039596, *Ae. aegypti* AAEL017774 and the consensus mosquito promoters performed consistently within each species, suggesting that these promoters might work in other mosquito species for which CRISPR reagents have not yet been optimized. Of the two *Drosophila* U6 promoters tested, only U6:3 resulted in significant KO effects in *Anopheles* and *Culex* cell lines but with very low efficiency (mean=4.3 SD±0.9; mean=7.4 SD±2). A secondary analysis of the flow cytometry data, performed using the variation of the median fluorescence intensity (MFI) of mCherry signal within the GFP^+^ cells, confirmed the relative changes in KO efficiencies (**Supplementary Figure 2b**). These results are in accordance with overall evolutionary distance between the species and U6 promoter sequence average distance and corroborate previous *in vitro*^10,11,13,14^ and *in vivo*^12,13^ results.

To facilitate CRISPR-based genome engineering in mosquitos and provide a batch-mode design resource for pooled CRISPR screening, we developed a new online resource for mosquitos, CRISPR GuideXpress (https://www.flyrnai.org/tools/fly2mosquito/web/),.which has a number of features. First, CRISPR GuideXpress allows users to input genes from *Drosophila*, which has a very well-annotated genome and is closely related to mosquitos, and automatically retrieves pre-computed sgRNA designs targeting the nearest ortholog in a selected mosquito species. Orthology is calculated using an approach similar to DIOPT^16^. Second, CRISPR GuideXpress facilitates search input and output in batch mode (i.e. for multiple genes at once). This allows simultaneous retrieval of large numbers of sgRNAs targeting multiple genes, as is required for CRISPR screen library design. Third, CRISPR GuideXpress supports several mosquito species— *An. gambiae, An. coluzzii, An. stephensi, C. quinquefasciatus, Ae. aegypti*, and *Ae. Albopictus—* and can be further updated to include additional species (**Fig. 2a**). Finally, the sgRNA designs are accompanied by pre-computed set of parameters that are displayed alongside sgRNA sequences and the total search output can be downloaded in table format. Efficiency predictions are calculated based on the ‘Housden score’^17^ and a machine learning-based analysis of *Drosophila* CRISPR cell screen data^9^. CRISPR GuideXpress also provides a cross-species reference when the same guide targets a homologous gene in one of the other species supported, allowing in some cases, inter-species targeting with the same reagents. For each mosquito species, the sgRNA designs cover ∼92-99% of protein-coding genes, and at least ∼62-93% of protein-coding genes are targeted by 6 or more high quality sgRNAs (i.e. designs with no predicted off targets). The number of designs and relative coverage per gene for mosquito genomes is similar to the library used for CRISPR KO screening in *Drosophila* cells^9,15^ (**Fig. 2b**,**c**). Furthermore, for *An. gambiae* and *An. coluzzii*, we incorporated a variant database based on full genome sequences of hundreds of field samples from the *Anopheles* 1000 Genomes Project^18,19^ in order to allow for selection of designs that would avoid common SNPs in wild populations (**Fig. 2d**).

**Fig. 2.**
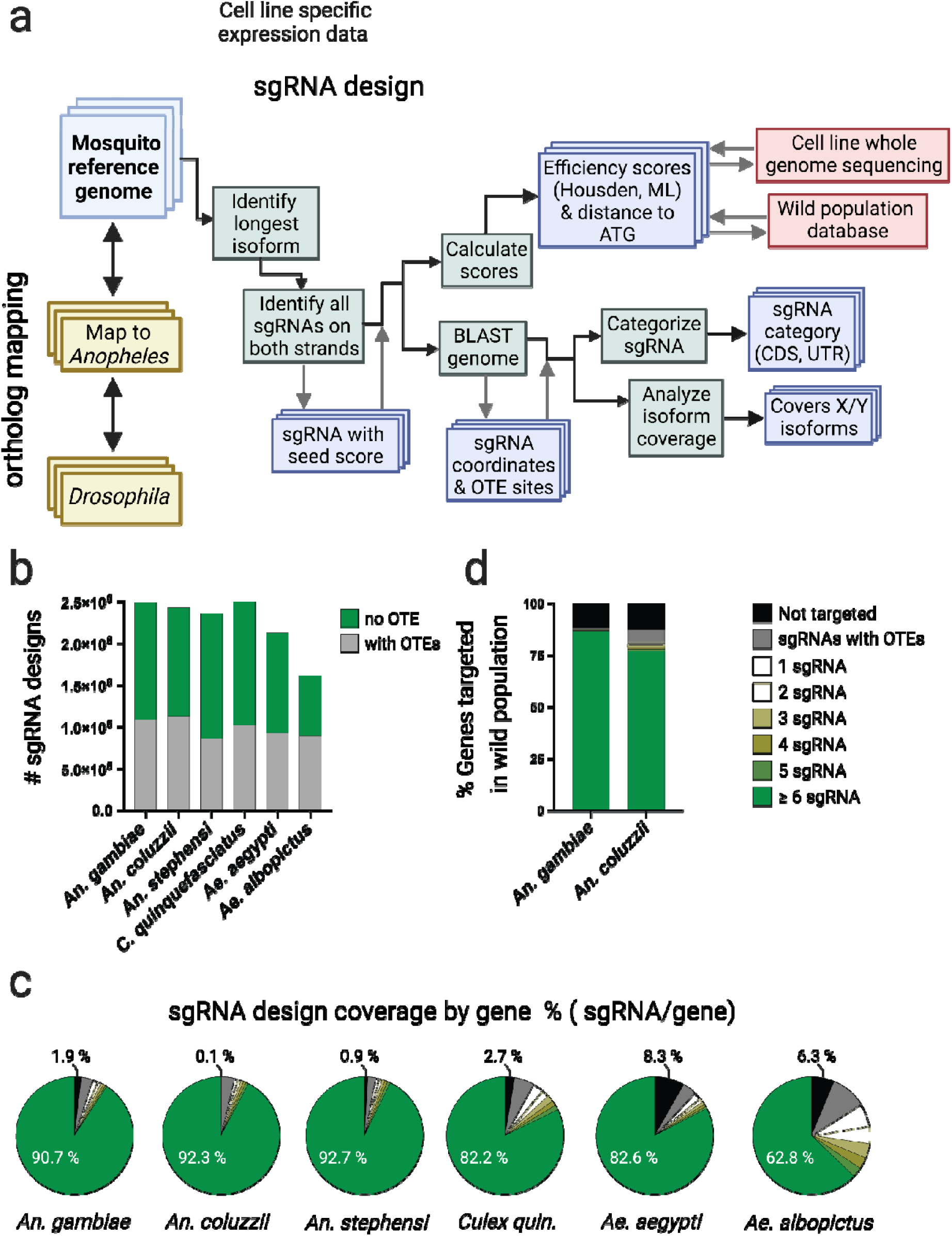
CRISPR GuideXpress: online bioinformatic framework for CRISPR sgRNA design and analysis. (**a**) Features and sgRNA design workflow. Ortholog mapping, cell line-specific expression data and sgRNA design for six supported mosquito species are integrated at one interface. Genes can be searched individually or in batch mode. Direct ortholog searching is available between *An. gambiae* and other mosquito species or *Drosophila*. After a gene name or ID is entered, the tool retrieves corresponding transcripts and displays precomputed sgRNAs and associated scores. The sgRNAs are computed as follows. The longest isoforms are identified from transcripts. Next, all possible PAMs and associated sgRNA designs on both strands are selected. Each design is then assigned a seed score based on uniqueness of the 12-15 nt 3’sequence (excluding the PAM). For each guide, a BLAST search is used to define specificity (off-target score). Each guide is mapped to the genome and categorized based on the gene region targeted and the respective isoform coverage. All sgRNA designs are evaluated to yield multiple efficiency parameters: ‘Housden’ score, machine learning (ML) score, and distance from ATG. Additionally, sgRNA designs for *An. gambiae* and *An. coluzzii* are assigned a ‘wild population efficiency’ score calculated from the Ag1000 Genome project dataset. To optimize for use in *An. coluzzii* Sua-5B cells, the tool indicates if the sgRNA sequences fully match the Sua-5B whole-genome sequence. **(b, c, d**) Analysis of genome-wide CRISPR KO sgRNA designs targeting protein-coding genes in supported mosquito species. (**b**) Histogram representing total number of sgRNA designs in two categories: 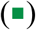 “no OTE” (off-target effect), with minimal off-target effects or 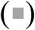 “with OTE” within the criteria (see Methods). (**c**) Genome-wide sgRNA design coverage, showing the percentage of genes targetable by sgRNAs with minimal OTE 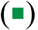, targetable only by sgRNAs with potential OTE 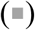, or untargetable (▪). (**d**) Genome-wide sgRNA design coverage by gene (%) in wild populations sampled in the Ag1000 Genome project. % of genes targeted and ranking based on # of sgRNAs/gene, as specified above. For this analysis were considered only sgRNA designs matching ≥ 95% of the wild genome sequences sampled.

Previous work in a *Drosophila* cell line showed that CRISPR screens can be conducted in an RMCE acceptor cell line by first introducing constitutive Cas9 expression and then transfecting cells with donor sgRNA expression vectors that can integrate into the RMCE locus^8^. Each cell stably integrates a small number of different sgRNA expression cassettes. Following outgrowth, sgRNAs that target essential genes are lost from the pool and following selection (e.g. with a toxic drug), sgRNAs that target genes for which knockout confers a growth advantage are enriched. To determine whether this approach can be used on mosquito cell lines, we used RMCE to integrate CRISPR sgRNAs into Sua-5B-1E8 cells stably expressing Cas9 (Sua5B-IE8-Act::Cas9-2A-Neo; **Fig. 3a**). Although we have authenticated these cells as derivative of *A. gambiae* M-form, now known as *Anopheles coluzzii*, we designed guides using gene sequences from the better annotated *Anopheles gambiae* genome (AgamP4, see Methods). For convenience, we refer to the *Anopheles gambiae* gene names throughout. We first confirmed that a visible phenotype can result from expressing an sgRNA expression cassette targeting *Rho1* (AGAP005160), which is necessary for the completion of cytokinesis. Previous reports have shown that knockdown of *Rho1* by RNAi in *Drosophila*^20^ or *Anopheles*^21^ cells results in a modest size increase (approximately 2-fold) due to cell growth without division, and *Drosophila* cells expressing CRISPR sgRNAs targeting *Rho1* become dramatically enlarged due to complete loss of *Rho1*^9^. To test the novel *Anopheles* cell-based CRISPR system, we transfected sgRNAs targeting the *Anopheles Rho1* ortholog AGAP005160 and observed transfected cells after several days of selection. We found that *Rho1* sgRNA-expressing cells, but not control cells, became enlarged up to 6-fold (**Fig. 3b**). We used T7 Endonuclease I assays to confirm editing of the *Rho1* locus (**Supplementary Figure 3a**). These results clearly demonstrate that using RMCE to deliver an sgRNA targeting an endogenous gene can result in a penetrant phenotype, suggesting that the system is compatible with CRISPR pooled-format screening.

**Fig. 3.**
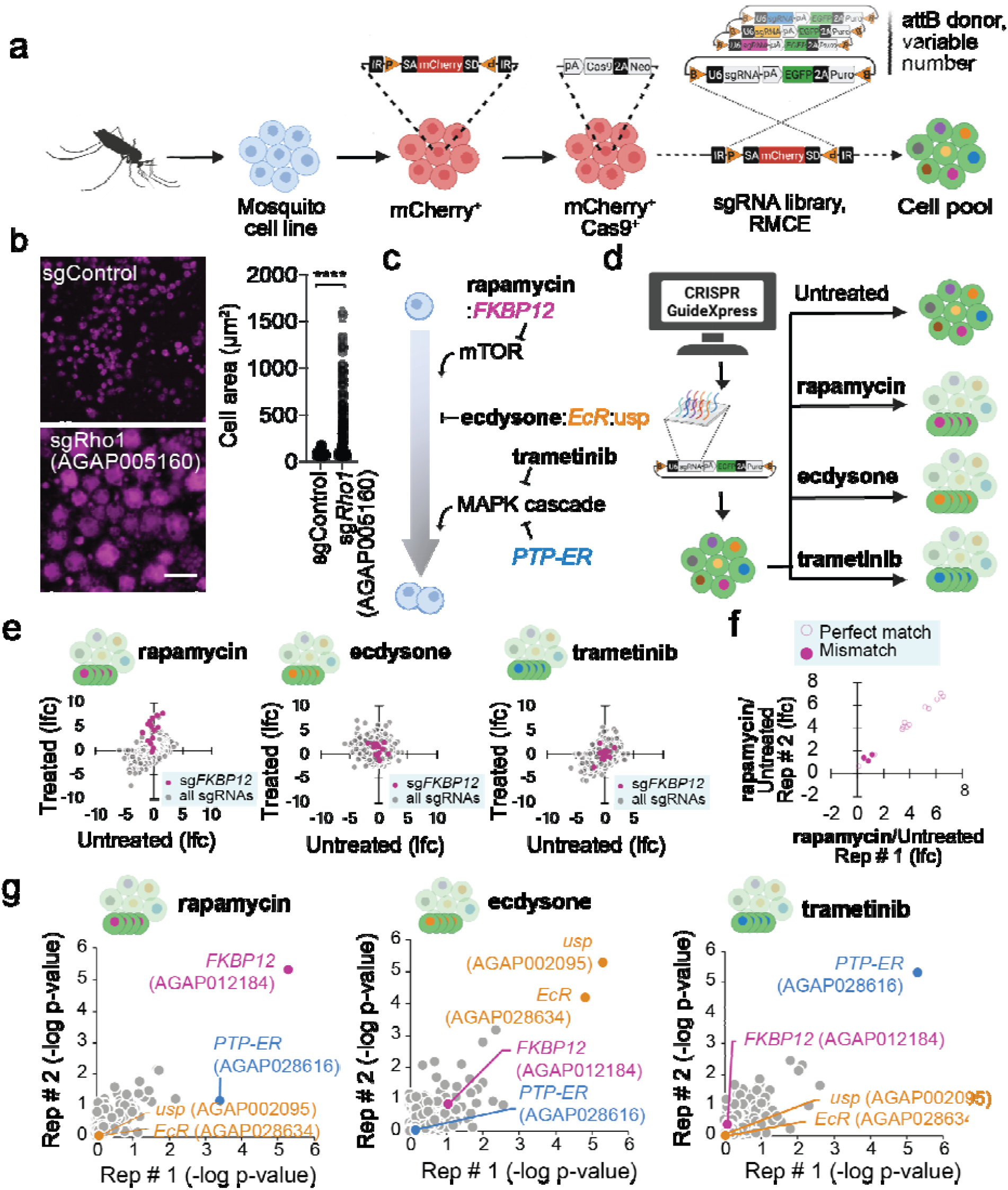
Pilot pooled CRISPR drug resistance screens in *Anopheles* cells. (**a**) Building CRISPR screen-ready cell lines. An *Anopheles coluzzii* Sua-5B RMCE acceptor cell line was stably transfected with Cas9 to create Sua5B-IE8-Act::Cas9-2A-Neo. Donor sgRNA vectors can now be used to create a screening pool. (**b**) Sua5B-IE8-Act::Cas9-2A-Neo cells were stably transfected with pLib6.4-Agam_695 donor vector encoding a *Rho1* sgRNA, leading to a highly penetrant failure in cytokinesis and dramatically enlarged cell area (****P<0.0001, unpaired t test, two-tailed, t=13.45, df=890, sgControl n=509, sgRho1 n=383).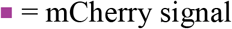, scale bar, 50 μm. (**c**) Schematic of proliferation-related pathways used to validate the screening approach. Rapamycin binding to FKBP12 inhibits mTOR, which is necessary for proliferation. Ecdysone binding to EcR and Usp inhibits proliferation. Trametinib inhibits the mitogen-activated protein kinase (MAPK) signaling cascade, which normally promotes proliferation. PTP-ER is a potent endogenous negative regulator of MAPK. (**d**) We used CRISPR GuideXpress to design a library of 3,487 sgRNAs against mosquito orthologs of *FKBP12, EcR, usp*, and *PTP-ER*. These were cloned into a library donor vector (pLib6.4-Agam-695) and integrated into Sua5B-IE8-Act::Cas9-2A-Neo cells. The cells were left untreated or treated with the indicated drugs. (**e**) Endpoint sgRNA readcounts compared to plasmid readcounts show that more than half of all *FKBP12* sgRNAs were enriched following growth in rapamycin but not ecdysone or trametinib. (**f**) FKBP12 sgRNAs predicted to target the reference genome, AgamP4, but with mismatches as compared to the Sua5B-IE8-Act::Cas9-2A-Neo genome failed to enrich following rapamycin treatment. (**g**) Robust rank aggregation (RRA) analysis of sgRNA readcount data from all screens shows that *FKBP12* was selectively enriched after rapamycin treatment, *EcR* and *usp* were selectively enriched after ecdysone treatment, and *PTP-ER* was selectively enriched in trametinib treatment (two biological replicates).

To directly test application of the CRISPR screening platform at large scale in mosquito cells, we used the *Anopheles coluzzii* Sua-5B-*IE8-Act::Cas9-2A-Neo* cell line to perform in parallel three proof-of-concept CRISPR knockout screens using a focused sgRNA library. We first chose five genes that had previously been shown to be drug-resistance factors in *Drosophila* cells^9,15^ and used CRISPR GuideXpress to design a library targeting their orthologs in *Anopheles coluzzi*. Target genes included *Anopheles* orthologs of *FKBP12* (AGAP012184), which encodes the cellular binding partner of the mTOR inhibitor rapamycin; *EcR* (AGAP028634) and *usp* (AGAP002095), which encode mediators of an antiproliferative transcriptional response to treatment with ecdysone; and *PTP-ER* (AGAP028616), which encodes a negative regulator of the mitogen-activated protein kinase (MAPK) signaling cascade that can be suppressed by treatment with the MEK inhibitor trametinib **(Fig. 3c; Supplementary Data 3)**. In total, 3,487 sgRNAs were synthesized and cloned into pLib6.4-Agam_695 containing the strong *Anopheles* U6 promoter and transfected into *An. coluzzii* Sua-5B-IE8-Act::Cas9-2A-Neo cells in the presence of ΦC31 integrase to facilitate recombination, then selected for 16 days in puromycin-containing media with continuous passaging every four days. A theoretical copy number of 1000 cells per sgRNA was maintained during all passages. For the selection screens, the cells were grown for an additional 30 days in the presence of rapamycin, ecdysone (20-hydroxyecdysone), or trametinib **(Fig. 3d**). Then, genomic DNA was collected and the sgRNA-containing locus was PCR amplified, barcoded, and analyzed by next-generation sequencing (NGS). Guides targeting *FKBP12* (AGAP012184) were clearly enriched by treatment with rapamycin but not in untreated, ecdysone, or trametinib growth conditions (**Fig. 3e**). Sequence analysis of the *FKBP12* locus in the Sua-5B-IE8-Act::Cas9-2A-Neo cell line revealed a coding variant in the cells relative to the reference genome (AgamP4) that results in single-base mismatches between a subset of three sgRNAs designed to target the *FKBP12* locus. Unlike no-mismatch guides, these mismatched guides were not selected in rapamycin treatment conditions (**Fig. 3f**). Similar observations were made for the set of guides targeting *usp* (**Supplementary Figure 3b**). After observing these single nucleotide polymorphisms in specific genes, we conducted whole-genome sequencing of the Sua-5B-IE8-Act::Cas9-2A-Neo ‘screen-ready’ cell line and added a variant analysis to CRISPR GuideXpress, giving users the option to exclude these variants from sgRNA designs (**Fig. 2a**). Importantly, for all three screens, we found significant and selective enrichment for the orthologs of the expected genes (i.e. *FKBP12* for rapamycin, *EcR* and *usp* for ecdysone, and *PTP-ER* in trametinib (**Fig 3g; Supplementary Figure 3c**). We note, however, that we did not observe enrichment for a candidate *EcI* ortholog. These results suggest that using the RMCE approach, optimized U6 expression, and CRISPR sgRNA design pipeline we have developed will make it possible to efficiently conduct massively parallel genetic screens in mosquito cells.

In conclusion, we have developed tools and methods that enable CRISPR pooled-format screens in mosquito cell lines that can now be used as a platform similar to those available in mammalian and *Drosophila* cells. This includes not only application of CRISPR-Cas9 knockout screening but also CRISPR/Cas-based activation and RNA knockdown screens. Moreover, CRISPR pooled screening in these cells can be combined with a variety of cell-based assays, including assays relevant to virus or parasite entry, innate immunity, resistance to insecticides and other toxins, and to investigate other health-relevant topics. We anticipate that data generated from such screens will substantially contribute to multifront efforts to control mosquito-borne diseases.

## Acknowledgments

We thank Flaminia Catteruccia and Nelson Lau for cell lines. We thank Daniela Silva-Ayala for helpful discussions. The research reported in this manuscript was supported by NIH NIGMS grant P41GM132087 to N.P. N.P. is an investigator of Howard Hughes Medical Institute.

## Author contributions

R.V and E.M. contributed equally to this work. R.V., E.M., Y.H., S.E.M. and N.P. conceived the project. R.V., E.M., Y.H., S.E.M, T.M.C. and N.P. contributed to the design of the experiments. J.R., P.M., and Y.H. developed the bioinformatic tools. R.V., E.M., and F.F.S. performed the experiments and contributed to the collection and analysis of data. E.M. wrote the first draft of the manuscript with input from R.V., S.E.M. and N.P. All authors edited and approved the manuscript.

## Data availability

The raw sequencing reads and VCF files of whole genome sequencing data from Sua-5B-IE8-Act::Cas9-2A-Neo cell line are available for download at CRISPR GuideExpress (https://www.flyrnai.org/tools/fly2mosquito/web/download).as well as the pilot CRISPR screen result.

