## Supplementary material for "Bioinformatic and cell-based tools for pooled CRISPR knockout screening in mosquitos": Supp Figures-Methods (from another version of this preprint: version 1, 2021-03-29)

Contents:

Supplementary Figure 1	3

Supplementary Figure 2	5

Supplementary Figure 3	6

Materials and Methods	7

Methods References	15

Additional Supplementary Data Online:

**Supplementary Data 1- List of primers, vectors and cell lines**

Includes a list of primers referenced in the Methods; all plasmids built or used for this study; parental and new engineered cell lines; sequences and features of U6 promoter expressing cassette used to build the attB library donor vectors. The data as described is subdivided in the following file tabs:

1. **Primers**
2. **Plasmids**
3. **Cell lines**
4. **Selected U6 promoter sequences:** Includes the sequence incorporated in the attB-library donor vectors and other structural details.

**Supplementary Data 2- Flow cytometry analysis data**

Data relevant to U6 promoter evaluation (data represented in **Fig.1c,d,e** and **Supplementary Figure 2b**). Includes output from FlowJo analysis and calculation of KO efficiency relative to the three mosquito cell lines analyzed. The datasets for each cell line are subdivided in a separate tabs:

1. **Sua-5B-IE8**
2. **Hsu-1.7**
3. **C6/36-HE8**

**Supplementary Data 3- Anopheles sgRNA Library and Readcounts**

Data relevant to *Anopheles* CRISPR screen (data represented in **Fig.3c-g** and **Supplementary Figure 3)**. Includes sgRNA library, readcount, analysis, and primers. The data as described is subdivided for each cell line in a separate tabs:

1. **sgRNA library**
2. **CRISPRseq Primers**
3. **Readcounts**
4. **Log2 fold-change**
5. **MaGeCK Test**
6. **Variant analysis**

**Supplementary Data 4- CRISPR GuideXpress sgRNA design statistics**

Includes raw statistics regarding sgRNA designs for all the supported species and data represented in **Fig.2b-e**. The data as described is subdivided in the following file tabs:

1. **Raw statistics**
2. **Data relative to Fig2b-d**

**Supplementary Figure 1**

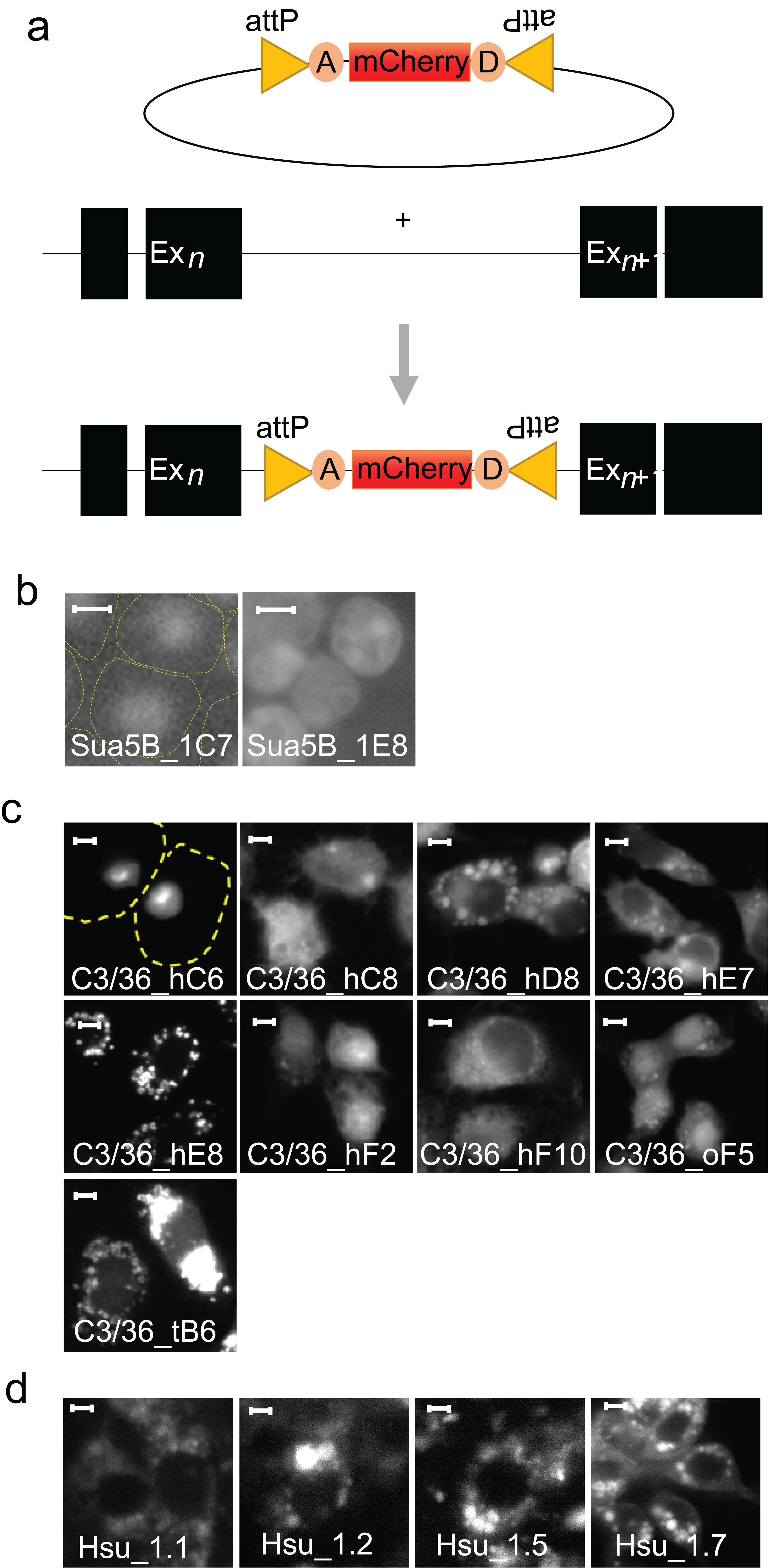

**Supplementary Figure 1**. **Building mosquito RMCE acceptor cell lines**. (**a**) A *Minos* transposon plasmid with promoterless mCherry was co-transfected along with a *Minos* transposase expression vector to generate intronic insertions that become artificial exons. In rare cases (~0.1%), this successfully results in expression of mCherry as a protein fusion. (**b-d**) mCherry positive *Anopheles* Sua-5B (b), *Aedes* C6/36 (c), or *Culex* Hsu (d) cells were isolated by fluorescence-activated cell sorting (FACS) and iteratively subcloned to obtain isogenic populations with bright mCherry signal. The mCherry positive cells were imaged, revealing a variety of intracellular distributions, consistent with exon-tagging of a variety of endogenous genes. Scale bar, 5 μm.

**Supplementary Figure 2**

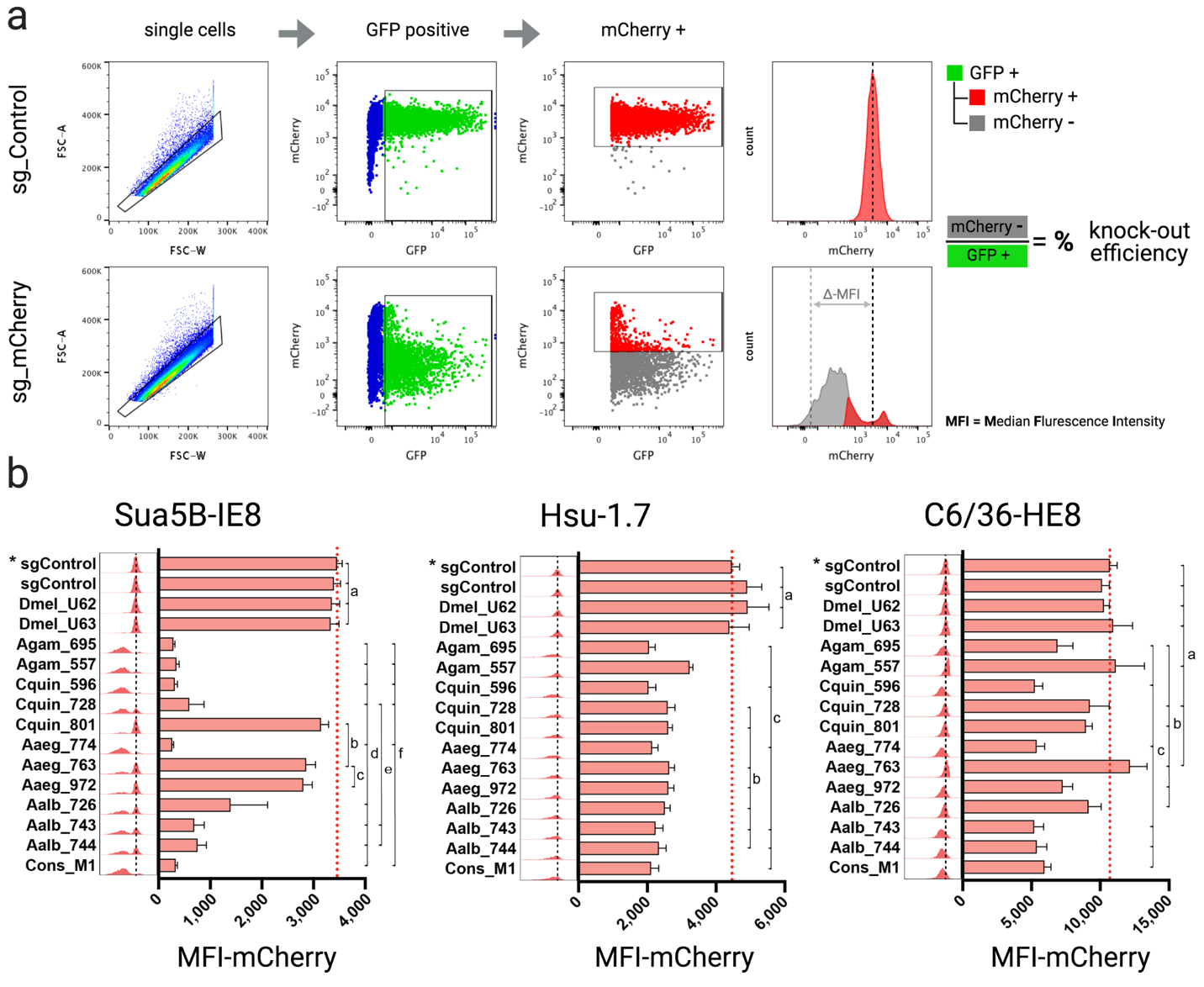

**Supplementary Figure 2**. **Flow cytometry gating strategy and supporting data on knockout efficiency with different U6 promoters**. (**a**) Schematic detailing the gating strategy used for flow cytometry analysis: single cells were gated based on forward scattering parameters; a subordinate gate was defined for GFP positive cells based on an un-transfected control; GFP positive population was sub-gated to define mCherry positive cells based on an mCherry-positive/GFP-positive control; GFP positive cells residing out of this gate were categorized as mCherry negative. The % of mCherry negative cells within the GFP population over the total GFP-positive population defines the knockout efficiency of the promoter tested. Alternatively, direct quantification of the MFI of the mCherry signal within the GFP-positive population and variation of MFI (ΔMFI) relative to control can be used to infer knockout efficiency. The MFI is a parameter of the population correlating directly with the number of mCherry molecules present in each cell, and thus can also be used to quantify KO of the mCherry fluorophore. MFIs are shown as dashed lines in the mCherry population histogram. (**b**) Histograms representing the MFIs of mCherry signals for each U6 promoter tested in the three cell lines. Bars represent the mean value of the MFI and error bars represent the SD from all replicates of three independent experiments. On the y-axis is also plotted a representative example of the mCherry signal fluorescence distribution within the GFP-positive population, with demarcation (dashed line) of the approximate position of the median in control samples. Statistical analysis was performed using Brown-Forsythe ANOVA test followed by Dunnett’s multiple comparison test. Lowercase letter groupings denote differences not significant (P_Dunnett_ > 0.05). All differences between samples of different groupings are significant (P_Dunnett_ < 0.05).

**Supplementary Figure 3**

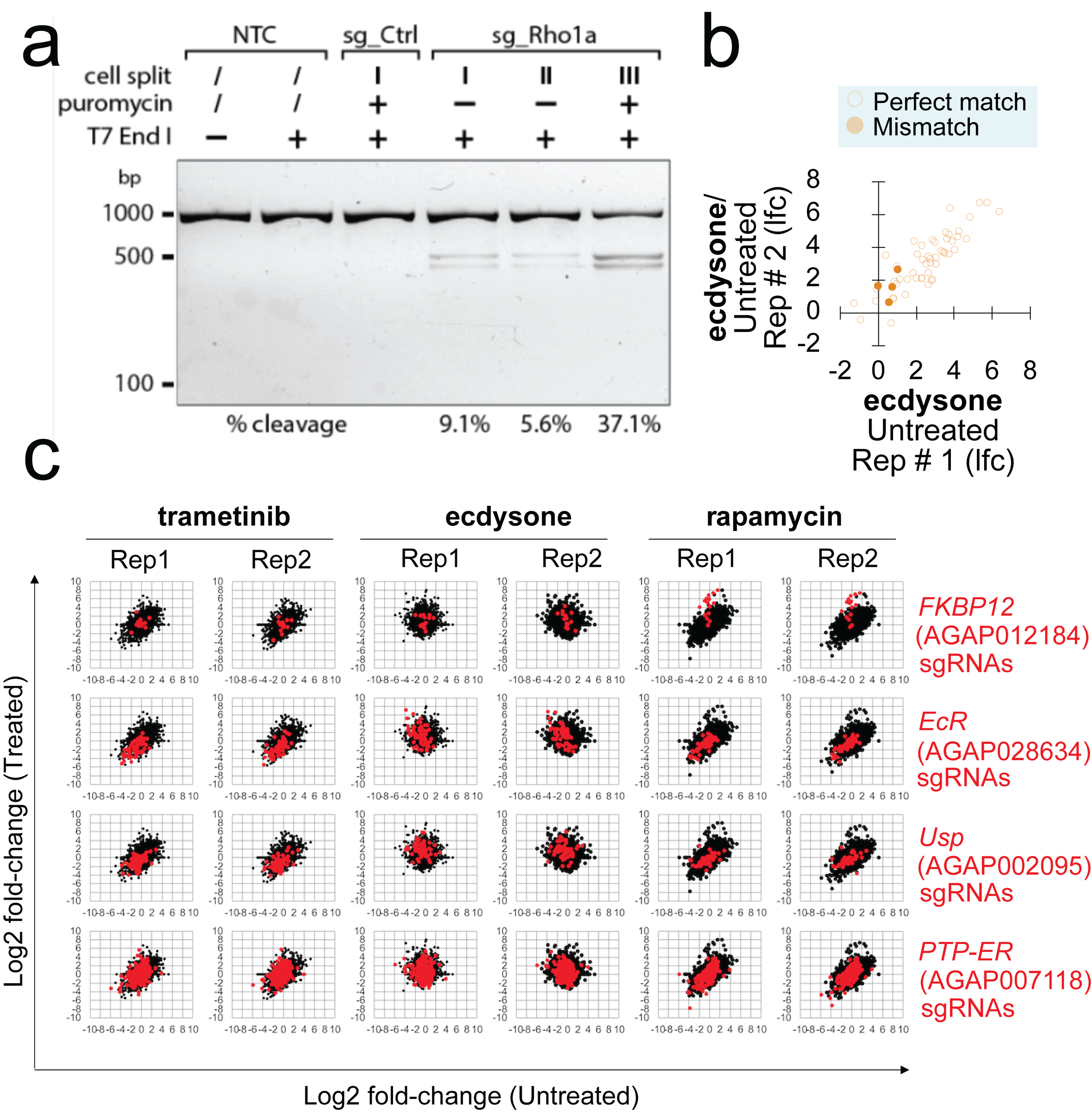

**Supplementary Figure 3**. **Further analysis of pilot *Anopheles* CRISPR screen data**. (**a**) T7 Endonuclease I assay confirming genomic editing in cells transiently transfected with Cas9 and a library vector with the highly efficient Agam_695 U6 promoter delivering either a non-targeting sgRNA as control (sg_Control) or a sgRNA targeting the gene that encodes the Rho1 GTPase (AGAP005160). (**b**) Enrichment of an sgRNA targeting *usp* (AGAP002095) in two biological replicates of the ecdysone screen. Variants in *usp* in Sua5B cells relative to the reference genome resulted in mismatches in 4 sgRNAs. Mismatched guides were not enriched following ecdysone treatment. (**c**) All enrichment data relative to plasmid library from pilot *Anopheles* CRISPR screens.

**Materials and Methods**

**Identification of mosquito U6 promoters**

U6 snRNAs are conserved eukaryotic non-coding RNAs that take part in the formation of the catalytic core of the spliceosome, while their promoters and associated *pol III* transcriptional machinery shows divergence even between closely related species. We used the *Drosophila* U6:2 snRNA sequence as query to perform a BLAST search of mosquito reference genomes, using the Vectorbase interface (now VEuPathDB at https://vectorbase.org/vectorbase/app).

Next, all sequences of the identified orthologs within each species were subjected to multiple alignment (using ClustalΩ at [www.ebi.ac.uk](http://www.ebi.ac.uk)^1^) including the full snRNA and 500 bp of upstream sequence, allowing us to visualize and exclude sequences lacking conserved portions of the pol III bipartite promoter motif (i.e. the PSEA and TATA). As a secondary criterion for selection, we relied on expression levels reported on RNA-seq data publicly available for each species, including tissue specific RNA-seq of adult mosquitos or cell lines (available on vectorbase/VEuPathDB). This data, although not fully reliable in reporting expression levels of small non-coding RNAs, was informative in narrowing our choice to putatively expressed snRNAs. Third, we consulted literature reports of the activity of these promoters.

After selecting up to three U6 promoters for each species, we performed a second inter-species alignment, and selected for each promoter a region upstream of the TSS of an arbitrary length ranging from 144-237 bp. In addition, we selected a 99 bp mosquito consensus sequence derived from the alignment. The length of the AGAP013695 promoter was chosen based on previous work from Konet et al.^2^ in cells and Hammond et al.^3^ in adult *Anopheles* mosquitos. The alignment shown in Fig.1a was obtained by aligning the 250 bp upstream of Drosophila U6-2 and U6-3 promoters to the selected mosquito sequences (only a 75 bp region is displayed). The mosquito consensus sequence is shown at the bottom. Jalview Version 2^4^ was used to visualize the alignment and to infer the phylogenetic hierarchy based on the average sequence distance of the snRNA and the 75 bp upstream sequence. Additional information about the U6 promoter sequences used for library vector construction is available in **Supplementary Data 1**.

**Cloning**

In order to build new library vectors expressing gRNAs under mosquito U6 promoters we synthesized ~500 bp gBlocks (Integrated DNA Technologies, Inc.) containing in order: the selected U6 promoter sequences; a BbsI cloning site (for gRNA insertion); the gRNA scaffold sequence including an at least 8-T termination sequence^5^; 30 nucleotides of the native termination sequence, and an additional portion of the termination sequence derived from the 3' end of the Drosophila snRNA-U6-96AB (CR32867). The gBlocks were sub-cloned into pCR™-Blunt II-TOPO® (Invitrogen) or directly digested with BstBI/KpnI and sub-cloned into pLib6.4 (EGFP reporter) or pLib6.4B (EBFP2 reporter) by replacing the entire gRNA expressing cassette. This resulted in 24 new library vectors harboring the mosquito gRNA expressing cassettes. An additional version of the pLib6.4 attB donor library vector, named pLib6.4B, was obtained by replacing EGFP with EBFP2 using overlap extension PCR with megaprimers as described in Bryksin et al.^6^ First, PCR was performed to amplify EBFP2 from pEBFP2-Nuc (Addgene #14893) using primers 14-15. Second, overlap extension PCR was performed using as megaprimers directly the EBFP2 amplicon to mutagenize the EGFP in pLib6.4. The same strategy, using the same primers, was also used to obtain pLib6.6B from pLib6.6, as well as EBFP2 and EGFP versions of the pSL1180-HR-PUbECFP plasmid (Addgene #47917). In order to generate the EGFP version, megaprimers were obtained by PCR amplification of EGFP from the pLib6.4 vector using the same primers (14-15).

Insertion of the gRNA sequence targeting mCherry or Rho GTPase in pLib vectors was performed following an established protocol^7^ that allows sgRNA oligos with BbsI sites to be annealed and then ligated into BbsI-digested pLib vectors (primers 1-13).

pDmAct5C::Cas9-2A-Neo plasmid was built from Ac5-STABLE1-neo (Addgene # 32425) by restriction/ligation cloning, replacing GFP with SpCas9, Drosophila codon-optimized from pl018^8^. pAaePUb::Cas9-2A-Neo was built in multiple steps. First, the Ae. aegypti polyubiquitin promoter^9^ (AAEL003888) was amplified from pSL1180-HR-PUbECFP (Addgene #47917) with primers 16-17 and assembled with EcoRV/SphI linearized pAW (Drosophila gateway vector collection) using Gibson assembly (New England Biolabs) resulting in pAePUbW destination vector, where the Drosophila Actin5C promoter was replaced with the *Aedes* polyubiquitin promoter. Second, the Gateway cassette in pAePUbW was removed by restriction with EcoRV/PmeI and replaced with Cas9-2A-Neo obtained with double restriction with BsaI/SalI from DmAct5C::Cas9-2A-Neo. When ligation-incompatible restriction extremities where generated, we performed blunting of 3’ overhangs using T4 DNA polymerase (New England Biolabs) and proceeded with ligation of blunt ends. A pActin-ΦC31-Integrase plasmid was obtained using the gateway recombination system. First the ΦC31-Integrase was amplified from pBS130 (Addgene #26290) using AttB tailed primers (18-19). Next, the AttB PCR-flanked product was used in a BP reaction to obtain a Gateway entry clone. The ΦC31-Integrase entry clone was used in a LR reaction with pAW destination vector (Drosophila gateway vector collection) to obtain the pA-ΦC3-Integrase expression vector. Cloning steps involving PCR amplification were performed using Q5 Hot Start High-Fidelity 2X Master Mix (New England Biolabs). All constructs were verified by targeted Sanger sequencing of the modified regions. All primers and plasmids used in this work are listed in **Supplementary Data 1**.

**Cell lines**

The *Anopheles coluzzii* cell line Sua-5B and *Culex quinquefascaitus* Hsu cell line were kind gifts from Flaminia Catteruccia (Harvard T.H. Chan School of Public Health, Boston MA) and Nelson Lau (Boston University, Boston MA), respectively. The *Aedes albopictus* C6/36 cell line was from the Colpitts lab. All cell lines were maintained at 25℃ in Schneider’s medium (Gibco), 1x Penicillin-Streptomycin (Gibco) and 10% heat inactivated fetal bovine serum (Gibco). In addition, the media for the Hsu cell line was supplemented with 1x MEM NEAA (Gibco). The *Anopheles coluzzii* Sua-5B cell line was authenticated by diagnostic PCR and variant calling analysis of cell-specific whole genome sequence data. Diagnostic PCR was performed following the protocol specified in Santolamazza et al.^10^ for *Anopheles* M/S molecular form discrimination (primers 20-21), confirming the presence of the SINE200 insertion specific to the M form (currently recognized as the *Anopheles coluzzii* species). Variant calling analysis (see “Genome variants” below) was performed by comparing whole-genome sequence data from the Sua-5B ‘CRISPR-ready’ cell line to *Anopheles gambiae* (AgamP4) or *Anopheles coluzzii* (AcolM1.8) genome sequence. We observed significantly more variants when the cell line sequence was compared with the *Anopheles gambiae* genome than when it was compared to *Anopheles coluzzii* (Agam variants=4,983,818; Acol variants=4,439,680; Δ=544,138).

**Cell line engineering**

Native cell lines were first transformed by transposition of an attP-flanked mCherry reporter cassette following an established protocol (**Fig. 3a; Supplementary Fig. 2a**)^11^. The reporter cassette containing both splice donor and splice acceptor sites, flanked by two inverted ΦC31 attP sites that allow for subsequent RMCE in the presence of the ΦC31-integrase and an attB-flanked donor cassette. Successful insertion through Minos transposition is revealed by mCherry fluorescence, as mCherry is spliced into the trapped gene as an artificial exon. This system offers the advantage to select for insertions in transcriptionally active regions of the genome, and acts as a docking pad for the delivery of complex sgRNA libraries through ΦC31 RMCE between attP sites in the cassette (genome) and attB sites of the library donor plasmid. Moreover, this system allows for monitoring of the rate of recombination by flow cytometry or microscopy as cells integrating the library cassette will lose mCherry fluorescence and report fluorescence from the exchanged library vector. Briefly, cells were co-transfected with Actin5C-Minos transposase vector and three separate MIMIC mCherry plasmids (encoding mCherry in each reading frame) in a single transfection mix (molar ratio 1:0.3:0.3:0.3) using Effectene (Qiagen) and following the manufacturer’s instructions for 12-well format. After transfection, cells were passaged two times, expanded, and single-cell sorted into 96-well plates by fluorescence-activated cell sorting (FACS) to obtain clonal mCherry-trapped populations. FACS was performed using a FACSAria or MoFlo Astrios (BD) with a 100 um nozzle at 20 psi (Harvard Medical School, Immunology Flow Core). For each cell line, multiple clonal populations were expanded and assessed based on signal intensity and subcellular localization of the mCherry reporter (example in **Supplementary Figure 1**). Selected Minos-transformed clones yielded the mCherry positive mosquito-derived cell lines we have named Sua-5B-IE8, Hsu-1.7, and C6/36-HE8, which were used for CRISPR KO efficiency dual-reporter assays of U6 promoter activity or further modified by stable transfection with Cas9 expressing plasmid. In particular, the *Anopheles* Sua-5b-IE8 clone was transfected with pAct::Cas9-2A-Neo and selected in 200 ug/ml Geneticin (Gibco) for 30 days to obtain our *Anopheles* CRISPR-ready cell line, Sua-5B-IE8-Act::Cas9-2A-Neo.

**U6 promoter evaluation**

CRISPR knockout efficiency of the stable mCherry reporter integrated in the mosquito cell lines was used as a readout to test relative U6 promoter sgRNA expression of different mosquito promoters, using transient transfection. Plasmid DNA for transfections was prepared using mini or midiprep kits (Qiagen) and quality analyzed by spectrophotometry and agarose gel electrophoresis. Plasmid DNA concentration was measured (Qubit, ThermoFisher) and plasmid mixes for transfection were normalized by copy number according to the M.W. of each plasmid and using pUC19 (ThermoFisher) to normalize total DNA amounts as needed. All cells were transfected with 300 ng of plasmid mix (12 well format) using Effectene (Qiagen) and following the manufacturer’s protocol. The % composition of each plasmid mixture used for transfection was as follows: for the *Anopheles* and *Culex* cell lines, a plasmid mixture containing 150 ng (50%) of pAct::Cas9-2A-Neo, and 150 ng (50%) of one of 14 different copy-number balanced mCherry-sgRNA expressing plasmids was transfected. Additionally, two control transfections were performed: one including the “empty” pLib6.4-Agam_695 expression vector (sgControl), which effectively drives the expression of a “non-targeting guide” matching the sequence of the empty BbsI cloning cassette, and a second including the same control guide but lacking the Cas9-expressing plasmid. Transfections in the *Aedes* cell line were performed with a different plasmid mixture containing 150 ng (50%) of pAaePUb::Cas9-2A-Neo as a source of Cas9 and 135 ng (45%) of one of 14 different copy-number balanced mCherry-sgRNA expressing plasmids and, in addition, 15 ng (5%) of pSL1180-HR-PUbEGFP-NLS to increase fluorescence output of the GFP reporter. For this cell line, we used the same sgControl plasmids, transfected with or without the Cas9-expressing plasmid. Additional information on plasmid vectors used is provided in **Supplementary Data 1**. Transfections were performed in 12-well format with 3 or more replicate wells per condition tested, and a total of three independent experiments were performed. *Culex* and *Ae. albopictus* cells were detached from flasks using Accumax (Innovative Cell Technologies, Inc) and seeded onto 12-well plates 16-24 h before transfection. *Anopheles* cells were resuspended from flasks by pipetting and seeded 30 minutes before transfection. *Culex* and *Aedes* cell lines were transfected at ~70% confluency, and *Anopheles* cells at ~90% confluency. 24 hours after transfection, cells were transferred to new plates and cultured for 12 days, then either analyzed by flow cytometry or slowly frozen at -80℃ in culture media supplemented with 10% DMSO (v/v) and stored until flow cytometry analysis. *Anopheles* and *Aedes* cells during the last passage preceding the experiment end point were treated with a sub-lethal dose of puromycin (0.5 ug/ml) to increase the ratio of transfected cells and reduce the volume of cells analyzed with flow cytometry (puromycin resistance is conferred by the sgRNA expression plasmid).

**Flow cytometry analysis**

Flow cytometry analysis was performed on a FACSymphony analyzer (BD) (Harvard Medical School, Immunology Flow Core). Immeditely prior to analysis, frozen cells were thawed quickly in a 30℃ water bath, washed twice in PBS, and resuspended in fresh culture media. Cells were loaded into 96-well plates and read using the high-throughput sampler. Flow speed, laser voltages and gating parameters were adjusted for each cell line. The gating strategy is shown in **Supplementary Fig. 2a**. Gates for cell singlets were defined based on forward and side scattering beams and a subordinate gate for GFP^+^ cells was defined based on untransfected control cells. GFP^+^ cells were sub-gated to define mCherry^+^ cells and mCherry^-^ cells within the GFP population were defined by exclusion from the mCherry^+^ gate (NOT Boolean gate for mCherry^+^). The CRISPR knockout efficiency relative to each U6 promoter tested was inferred using the populations defined above as: knockout efficiency % = {[# of mCherry^-^ cells] ÷ [# of total GFP-positive cells]}*100. We also measured the median fluorescence intensity (MFI) of the mCherry signal within the GFP population as an alternative method to quantify KO of the mCherry reporter. MFI is a parameter of the population that correlates with the number of molecules present in each cell, and thus can also be used to quantify KO of a fluorescent reporter.

All analysis were performed using FlowJo 10.7.1 (BD). To adjust for differences in the number of cells analyzed for each sample within a replicate experiment, cells defined by the GFP gate were down-sampled to a fixed arbitrary number using the DownSample plugin. Samples with <1000 GFP-positive cells were discarded from the analysis.

**Rho 1 GTPase targeting**

To target the Rho1 GTPase locus (AGAP005160) in Sua5B-IE8-Act::Cas9-2A-Neo cells, we seeded approximately 1.8×106 cells in 12-well dishes for 30 minutes and transfected using Effectene (Qiagen) and following the manufacturer’s instructions. Cells were transfected with a plasmid mixture containing 150 ng (50%) of pActin-ΦC31-Integrase and 150 ng (50%) of either pLib6.4-Agam_695 “empty” (sg_Ctrl) or a sgRNA targeting Rho1, and then cultured for 7 days in 4.5 μg/ml puromycin selective media. Cells were transferred to 96-well plates (Falcon #353219) and images of control or Rho1 sgRNA-treated cells were captured with a IN Cell Analyzer 6000 confocal imaging system (GE) using a 20X objective and dsRed filter setting to record mCherry signal stably reported from the cell line. Measurements of cell area for quantification of the Rho1 phenotype were performed using one field of each control or Rho1 sgRNA-treated cells using Fiji image analysis software. Briefly, images were converted into 8-bit binary images; thresholding based on the mCherry signal was performed using the “Li” algorithm, followed by “fill holes” and “watershed” processing algorithms, then the number of cells and associated surface area were calculated.

We used the T7-endonuclease I assay to verify editing at the Rho1 gRNA targeted region. First, genomic DNA was isolated using a Quick DNA Miniprep Kit (Zymo Research) according to the manufacturer’s instructions. Phusion polymerase (New England Biolabs) using HF buffer with primers 22-23 was used to obtain a PCR amplicon (877 bp) asymmetrically spanning the genomic target site of the sgRNA on the Rho1 GTPase locus (AGAP005160). The PCR reaction was directly denatured at 95℃ for 5’ and slowly cooled down to room temperature to allow re-annealing. Next, 8 ul of the resulting PCR reaction was combined with 1 μl NEB2 buffer and 1 μl T7 Endonuclease I (New England Biolabs) , digested 15’ at 37℃, then resolved on a 3% TBE agarose gel at 160 V for 1.5 hours. A gel image was acquired using a Bio-Rad ChemiDoc imaging system and relative cleavage % was inferred by determining band intensities using Fiji image analysis software and dividing the intensity of digested amplicon to that of the undigested amplicon in each lane.

**Library design, and cloning**

sgRNAs were chosen using CRISPR GuideXpress. Briefly, a list of five *Drosophila* genes (*FKBP12*, *EcR*, *usp*, *Oatp74D*, and *PTP-ER*) were inputted and all computed sgRNAs were retrieved. Additionally, a second set of *Drosophila* genes from a previous study^12^ was inputted, and the top six sgRNAs per gene were selected based on the following criteria: minimal OTE (off-target effect) score, maximum ML (machine learning efficiency) score BbsI-site bearing sgRNAs were culled from batch results. sgRNA sequences were placed within a 109-mer using specific tags and retrieved using dial-out PCR^13^. Each dial-out product was digested with BbsI and separated on a 20% non-denaturing polyacrylamide TBE gel (ThermoFisher Scientific). The product was next extracted using the crush-soak method and ligated into BbsI-digested pLib6.4-Agam_695. The product was then transformed into E. cloni 10GF' ELITE Electrocompetent Cells (Lucigen) and plated onto ten 15 cm LB plates containing carbenicillin and grown overnight at 30°C. A total of 100 bacterial colonies per construct were harvested into 25 mL of LB medium, mixed with an equal volume of 50% glycerol and stored in 1 mL aliquots at -80°C. Prior to transfection, the plasmid library was prepared by miniprep (Qiagen).

**Chemical-genetic CRISPR screening**

Sua5B-IE8-Act::Cas9-2A-Neo cells in log phase of growth were seeded at 6 ×10^6^ cells per 6-well dish in growth media containing antibiotics. They were transfected with a plasmid mixture containing equimolar amounts of pActin-ΦC31-Integrase and sgRNA donor plasmid library (pLib6.4-Agam_695) using Effectene (Qiagen) according to the manufacturer’s base protocol (“1:25”). Transfection efficiency was found to be 22-28% under these conditions. To achieve 1000 cells per sgRNA, 3,487 sgRNAs × 1000 cells/sgRNA ÷ 0.2 = ~18 ×10^6^ were transfected in three wells of a 6-well dish. After 4 days, each well was expanded into a 100 cm dish containing 5 μg/mL puromycin. Cells were cultured for an additional 12 days with media changes every 4 days. This achieved an integration percentage of approximately 70%. Cells were transferred to media containing puromycin and selective drugs, determined to be around the IC50 for Sua5B-IE8-Act::Cas9-2A-Neo cells: rapamycin (LC Laboratories) was dissolved in DMSO and used at a final concentration of 40 nM; 20-hydroxecdysone (Sigma) was dissolved in ethanol and used at a final concentration of 50 ng/mL; trametinib (Sellek Chemical) was dissolved in DMSO and used at a final concentration of 400 nM. Cells were continually passaged in selective medium for 30 days with media changes or re-seeding. Re-seeding density was maintained above 1000 cells/sgRNA at all times. Following selection, cell pellets representing >1000 cells/sgRNA were extracted using the Quick-gDNA MiniPrep kit (Zymo). Next, the genomic DNA was subjected to 2-step PCR to introduce in-line barcodes, a variable fingerprint, and Illumina sequencing primer and adapters. Amplicons were subjected to sequencing using a NextSeq500 at the Harvard Biopolymers Facility at Harvard Medical School. Computational barcode removal was performed using in-house scripts. Low-read sgRNAs (those with fewer than 10 reads in the plasmid library) were removed from the readcount files. All subsequent readcount and data analysis was performed using MaGeCK 0.5.7.

**Orthology mapping, sgRNA design, and variant analysis**

*Ortholog mapping between Drosophila melanogaster* (Dmel) *and Anopheles gambiae* (Agam). Ortholog mapping was extracted from the following five different prediction algorithms: orthoMCL vs5, eggNOG vs5.0, InParanoid vs8, orthoFinder (for which the code was run locally using the protein reference of Agam from VectorBase and Dmel from RefSeq), and TreeFam vs9. Orthologous pairs were integrated using the same pipeline as DIOPT^14^ and each orthologous pair was assigned a DIOPT score, i.e. the count of integrated algorithms that predict the pair (maximum score of 5).

*Ortholog mapping among mosquito species*. Ortholog mapping was obtained using Biomart at Vectorbase (https://biomart.vectorbase.org/biomart/martview). The Biomart ortholog mapping used for this study was subsequently replaced when Vectorbase merged with EuPathDB to become VEuPathDB. Ortholog mapping is now available from the new user interface using a gene ID search strategy with an ortholog transform step (https://vectorbase.org/vectorbase/app/query-grid).

*sgRNA design pipeline.* Supported species and the information about reference genome versions are available at the dropdown menu on the CRISPR GuideXpress search page (https://www.flyrnai.org/tools/fly2mosquito/web/species). The species currently supported at CRISPR GuideXpress and their corresponding genome versions are as follows: *Anopheles gambiae* (AgamP4.12), *Anopheles coluzzii* (AcolM1.8), *Anopheles stephensi* (AsteS1.7), *Aedes aegypti* (AaegL5.2), *Aedes albopictus* (AaloF1.2 & C6/36), and *Culex quinquefasciatus* (CpipJ2.4). Genome sequence and annotation files were obtained from https://vectorbase.org/vectorbase/app/downloads/Pre-VEuPathDB%20VectorBase%20files/. Input files used by the pipeline include

<species_name>_BASEFEATURES_<species_version>.gtf, <species_name>_TRANSCRIPTS_<species_version>.fa, and <species_name>_CHROMOSOMES_<species_version>.fa.

The pipeline starts by using the genome annotation to determine the transcript with the longest coding sequence (CDS) for each gene. In cases where there is a tie, the first transcript is chosen arbitrarily. Next, all potential sgRNA designs within the CDS of the selected transcripts are identified and logged. Then, unique k-mers within the genome are identified for computing seed scores for each design. Design sequences are BLASTed against the genome sequence and off-target scores are assigned based on off-target alignment hits with varying numbers of mismatched nucleotides. Off-target hits with fewer mismatches are weighted more heavily. The OTE (Off-target effect) score was calculated based on the number of potential off-target sites at 3 different thresholds and defined as follows: “OTE score= a.bc” where (a) is added to the digit before decimal point and is the number of off-target sites of the least stringent threshold (only considering off-target sites with 3 or fewer mismatches); (b) is added to the digit after the decimal point and is the number of off-target sites at the moderate threshold (only considering off-target sites with 4 or fewer mismatches) (c) is added as second digit after the decimal place and is the number of off-target sites under the most stringent threshold (5 or fewer mismatches). The categorization of sgRNA designs in Fig.2 b, c, e was defined as follows: “no OTE” (least stringent criteria)= score <1; with OTE (least stringent criteria) = score ≥ 1.

For each sgRNA, a Housden efficiency score^8^ is computed using a position matrix, and a machine learning-based efficiency score is computed using the pipeline available here: https://github.com/PierreMkt/Dmel-sgRNA-Efficiency-Prediction, which is based on *Drosophila* cell CRISPR screens^15^. Finally, a comparison with SNPs associated with genome data from wild populations is calculated based on the Ag1000G dataset^16^: https://www.malariagen.net/data/ag1000g-phase-2-ar1 (available for AgamP4.12 and AcolM1.8). The sgRNA pipeline was written in Python 3 and Perl 5.24.0, and uses BLAST 2.6.0. See **Supplementary File 4** for sgRNA design statistics.

*Analysis of wild populations variants*. Efficiency of sgRNAs for *An. gambiae* and *An. coluzzii* in wild populations was evaluated based on the datasets from *Anopheles gambiae* 1000 genomes project^16,17^ (Ag1000G) (https://www.malariagen.net/data/ag1000g-phase-2-ar1). Percent efficiency in wild populations was calculated based on the percent of samples carrying the SNP in each of the sgRNA target sequence. The SNP analysis results were obtained from <ftp://ngs.sanger.ac.uk/production/ag1000g/phase2/cas9_targets/> ^18^while the sample metadata was obtained from <ftp://ngs.sanger.ac.uk/production/ag1000g/phase2/AR1/samples/>^16^

*Genome variants in Sua-5B cell line.* The Sua-5B cell line was sequenced by BGI Genomics (<https://www.bgi.com/global/>) and processed as follows. The paired-end sequencing data was QC'd using fastqc 0.11.8 and multiqc 1.9. Pre-processing was performed according to the Broad Institute's best practices workflow for data preprocessing for variant discovery (https://gatk.broadinstitute.org/hc/en-us/articles/360035535912-Data-pre-processing-for-variant-discovery). First, the raw reads were aligned to the *Anopheles gambiae* (AgamP4) genome sequence or separately, to the *Anopheles coluzzii* (AcolM1.8) genome sequence using the Burrows-Wheeler Aligner BWA-MEM algorithm (http://bio-bwa.sourceforge.net/). Next, MarkDuplicatesSpark from the Broad's Genome Analysis Toolkit Genome (GATK) was used to identify read pairs that likely originated from the same original DNA fragments due to artifacts (https://gatk.broadinstitute.org/hc/en-us/articles/360050814112-MarkDuplicatesSpark). Variants were identified by following the Broad Institute's best practices workflow for germline short variant discovery (https://gatk.broadinstitute.org/hc/en-us/articles/360035535932-Germline-short-variant-discovery-SNPs-Indels-). Variant calling was performed using the GATK HaplotypeCaller (https://gatk.broadinstitute.org/hc/en-us/articles/360037225632-HaplotypeCaller). Finally, hard filtering was performed in accordance with the Broad Institute's recommendations for generic hard filtering (https://gatk.broadinstitute.org/hc/en-us/articles/360035890471-Hard-filtering-germline-short-variants). GATK version 4.1.8.1 was used for the analysis.

To obtain RNA-seq expression data from Sua-5B cells, the raw sequencing files were first obtained from VectorBase (<https://vectorbase.org/vectorbase/app/record/dataset/DS_1c16f776df>), QC'd using fastqc 0.11.8 and multiqc 1.9, and analyzed using Salmon 1.3.0 (https://combine-lab.github.io/salmon/) and tximport 1.16.0 (https://github.com/mikelove/tximport).

**Implementation and user interface (UI) of CRISPR GuideXpress**

*Implementation*. CRISPR GuideXpress is available at https://www.flyrnai.org/tools/fly2mosquito. The backend is written in PHP using the Symfony framework and precomputed results are stored in a MySQL database. The frontend uses the Twig template engine, Bootstrap, and some custom CSS for the UI. The JBrowse genome browser is used to display results visually and JQuery with a DataTables plugin is used for result tables. The website is hosted by the Research Computing group at Harvard Medical School.

*UI features.* The UI provides three major functions. First, the resource supports ortholog mapping of lists genes between *Drosophila* and *An. gambiae* and among supported mosquito species, including a ranking when one input gene maps to multiple genes. Second, the resource supports searches for sgRNA designs for a single gene or a list of genes. Both specificity scores, including the seed score and OTE score, and predicted efficiency scores are provided, allowing users to select optimal designs from multiple target sites. In addition, genome variants in *Anopheles coluzzii* Sua-5B cells were annotated for each sgRNA design, so that users can avoid target sites that might fail to work in this cell line. Each sgRNA design is also annotated with regards to SNPs as compared with the Ag1000 wild population genome data. The resource also supports search of expression levels in the *Anopheles coluzzii* Sua-5B cell line, starting with a single gene or a list of genes.

**Additional software**

Figures and graphical elements in this manuscript were created and assembled with BioRender and Adobe Illustrator. Statistical analysis were performed and plots were drawn with GraphPad Prism 8.

**Methods references**
